# From Brown to White: Brown Adipose Tissue Endothelial Cells whiten in Culture Conditions

**DOI:** 10.1101/2025.09.17.676510

**Authors:** Tabea Elschner, Staffan Hildebrand, Jana Sander, Lara Heubach, Nina Pannwitz, Stephan Grein, Maria Mircea, Elba Raimundez, Anastasia Georgiadi, Joerg Heeren, Jan Hasenauer, Alexander Pfeifer, Kerstin Wilhelm-Jüngling

## Abstract

Endothelial cells (ECs) are central regulators of vascular and metabolic homeostasis, yet their organ- and depot-specific diversity remains underexplored. Two major types of adipose tissue (AT) can be distinguished that differ substantially in their physiological function and vascularization: white AT (WAT), which is the major energy storage and brown AT (BAT), which is highly vascularized and dissipates energy [1–5]. While ECs from these depots likely contribute to adipose function, their characterization has been hindered by technical limitations in isolation and culture. Here, we establish a protocol for isolating and expanding ECs from murine BAT and WAT, enabling transcriptomic and functional analyses across depots. We demonstrate that freshly isolated BAT-ECs express depot-specific gene signatures, including *Rgcc*, *Cdkn1c*, *Tcf15*, *Meox2*, and *Efnb1*, several of which are dynamically regulated during cold-induced BAT activation. These findings reveal novel BAT-EC markers and highlight specialized endothelial programs that may support thermogenesis. However, we also uncover that culturing BAT-ECs profoundly remodels EC identity. Transcriptomic profiling shows that BAT-ECs rapidly downregulate BAT-specific markers and acquire features resembling WAT- and lymphatic ECs. This dedifferentiation is accompanied by signatures of proliferation, adhesion remodeling, and endothelial-to-mesenchymal transition. While these changes present challenges for maintaining depot-specific identity in culture, they also provide a framework to better interpret experimental outcomes and to investigate EC plasticity. Taken together, our study delivers a novel isolation and culture protocol for adipose ECs, defines BAT-EC markers, and demonstrates how culture conditions reshape their identity. These insights build the foundation for future research of AT vasculature.

**Highlights:**

- Fast method to purify endothelial cells (ECs) for direct RNA or protein extraction.
- Easy method to isolate and culture ECs specifically from thermogenic adipose tissue.
- Novel signature of BAT-ECs regulated upon browning and lost in culture.
- RGC-32, p57^kip2^, TCF15/MEOX2 and EFNB1 as novel BAT-EC-specific markers.

## 1. Introduction

Endothelial cells (ECs) line the inner surface of blood vessels and are essential regulators of tissue homeostasis, angiogenesis, and immune cell trafficking. Once thought to be relatively uniform, it is now well established that ECs exhibit a high degree of heterogeneity across organs and vascular beds [6–8]. Single-cell transcriptomic studies have demonstrated that ECs acquire organ-specific gene expression profiles in response to local cues such as shear stress, oxygen tension, growth factors, and interactions with surrounding parenchymal cells [6,9]. This endothelial diversity is not only structural but also functional, as ECs integrate tissue-specific signaling networks and actively participate in organ physiology and pathology [7].

*In vitro* culture of ECs has long been a valuable tool in vascular biology, providing a simplified and controllable system for studying endothelial function, angiogenic signaling, and vascular inflammation. Human umbilical vein endothelial cells (HUVECs), human dermal microvascular ECs (HDMECs), and immortalized lines such as EA.hy926 are among the most commonly used models. These cells have yielded key insights into EC biology and remain widely used due to their accessibility and ease of manipulation. However, it is increasingly clear that such generalized models may fail to recapitulate the unique characteristics of organ-specific ECs [8]. For example, ECs from the brain express tight junction proteins and transporters absent in other vascular beds [10], while liver sinusoidal ECs are fenestrated and display distinct scavenging functions [11]. Recently, also ECs of adipose tissues have been shown to vary between depots and to be highly influenced by the metabolic status of the organism [12–14]. The culture environment inevitably imposes adaptive changes on EC phenotype. When removed from their native microenvironment, ECs are forced to proliferate and therefore undergo a certain degree of remodeling. This includes the loss of *in vivo* cues such as laminar shear stress, extracellular matrix interactions, and intercellular communication, which together drive cells to adjust their transcriptome and growth behavior [15]. Rather than viewing this as a limitation, we propose that such remodeling reflects a necessary step for ECs to exit their highly specialized *in vivo* state and enable robust proliferation *in vitro*.

In metabolic research, the role of ECs is gaining increasing attention, especially adipose tissue (AT) EC. Importantly, not all adipose depots are the same: white adipose tissue (WAT) primarily stores energy, while brown adipose tissue (BAT) has thermogenic function and dissipates energy through mitochondrial uncoupling. In addition, to these major forms of AT, an intermediate form has been described, so called “beige fat” with the inguinal subcutaneous depot (WATi) containing the highest number of beige adipocytes, especially after cold exposure [1–5]. These differences are mirrored at the level of ECs. AT-ECs have been shown to regulate lipid transport, immune cell infiltration, and thermogenic programming [12,16,17]. Yet, despite their importance, depot-specific ECs remain under-characterized, largely due to challenges in their isolation and limited access to reliable culture systems. Therefore, most mechanistic insights are derived from non-depot-specific or immortalized cells. This raises critical questions: Can we develop a robust method to isolate and expand ECs from WAT and BAT? If so, do these cells exhibit a depot-specific identity *in vitro*, or do they converge toward a common, culture-induced phenotype?

In this study, we aimed to address these questions by establishing a robust protocol for the isolation and culture of adipose-derived ECs. Our method enables transcriptomic profiling and functional characterization of ECs from distinct adipose depots, while also uncovering the extent to which culture conditions influence their identity. Understanding how *in vitro* conditions shape adipose EC phenotypes is essential for interpreting data generated from culture models—and may also provide insight into how ECs respond to metabolic stress *in vivo*.

## 2. Experimental procedures

### 2.1. Animals

To establish the EC isolation protocol, mice of a C57BL6/J genetic background were used. For the isolation from pups, litters of early neonates were used, pooling 3-5 animals per isolation batch. For the isolation from adult mice, 8-16 week old mice were harvested and endothelial cells isolated from single mice. Both male and female mice were used. All animal experiments have been approved by the local authorities including the Committee for Animal Rights Protection of the State of North Rhine Westphalia (Landesamt für Natur, Umwelt und Verbraucherschutz: Az81-02.04.2022.A182).

### 2.2. Cold acclimation

C57BL6/J animals were kept in individually ventilated cages in a specific pathogen-free animal facility with controlled light–dark cycle (12 h–12 h), humidity (50-70%) and temperature (4-23°C). Mice were housed individually with reduced cage bedding and allowed to acclimatise to the housing conditions for 48 h. After that, housing temperatures were decreased to 16°C for 72 h to allow for cold acclimatisation. Then, housing temperatures were decreased to 4°C for seven days. Control animals were housed at 23°C for twelve days. Mice had access to food and water *ad libitum* throughout the experiment.

### 2.3. EC isolation

For EC isolation from pups, interscapular BAT and inguinal WAT of early neonates was dissected after decapitation. Adipose tissues of 3-5 neonates were pooled and processed one batch in the EC isolation protocol. Adult mice were killed by cervical dislocation and interscapular BAT, inguinal WAT and gonadal WAT were dissected. Tissues of adult mice were processed individually or pooled for up to three animals. For EC isolation after chronic cold exposure, adipose tissue of single mice was processed. Tissues were cut into small pieces and placed into adipose tissue dissociation mix in gentleMACS™ C tubes (Miltenyi Biotec, 130-0930237). For one batch of adipose tissue dissociation mix, 1.25 mL DMEM (gibco, 11995-065) plus 50 µL Enzyme D, 25 µL Enzyme R and 6.25 µL Enzyme A (Miltenyi Biotec, 130-105-808) were used. Tissues were dissociated on gentleMACS™ Octo dissociator with heaters on “37C_mr_ATDK_1” setting. The cell suspension was filtered using 100 µm and 40 µm cell strainers, washing both the collection tubes and strainer with sterile 0.5% bovine serum albumin in PBS (PBS-B). The cell suspension was washed by centrifugation 300 xg 5 min at room temperature and resuspension in 10 mL PBS-B. Through two consecutive centrifugation steps, the cell suspension was concentrated down to first 1 mL and then 90 µL cell suspension in PBS-B. Cells were incubated for 10 min on ice with FcR blocking reagent (Miltenyi Biotec, 130-092-575) to block unspecific binding.

The day before the intended EC isolation, Dynabeads™ (Thermo Fisher Scientific, 11035) were prepared for overnight coupling to CD31 antibody (BD Pharmigen, 553370). For this, 18 µL Dynabeads™ were washed with PBS-B on a magnetic separation rack (Cell Signaling Technology, 14654S) and resuspended in in 296,25 µL PBS-B and 3,75 µL CD31 antibody in an 1,5 mL tube. The tube was mounted on an overhead rotator and rotated overnight at 4 °C. On the day of the experiment, the Dynabeads™ were washed in PBS-B to remove unbound antibody and resuspended in 300 µL antibody. After FcR blocking, the tissue samples were resuspended with 50 µL PBS-B and 50 µL washed antibody-coupled Dynabeads™. The sample tube was mounted on an overhead rotator and rotated for 1 h at room temperature. Using a magnetic separation rack, the bead-labelled cells were washed before resuspension in endothelial cell medium. Endothelial cell medium consisted of DMEM F12 (gibco, 21041-025), 20% fetal calf serum (gibco, 10270-160), 4 mL Endothelial Cell Growth Supplement / Heparin (ECGSH, PromoCell, C-30140), 1 % Penicillin/ Streptomycin (P 10,000 units/mL, S 10,000 µg/mL, gibco, 15140122) and Amphotericin B (0,5 µg/mL, Carl Roth, 0246.1). For cultivation, the isolated ECs were seeded in 12 well plates at about 200.000 cells/well. The wells were pre-coated with 1% gelatin in H_2_O for 1 h at 37°C. For neonate isolations, the protocol yielded on average 305 000 (± 95 000) BAT-ECs and 470 000 (± 260 000) WATi-ECs, for EC isolation from adult mice on average 120 000 (± 90 000) BAT-ECs, 1 530 000 (± 350 000) WATi-ECs and 1 430 000 (± 470 000) WATg-ECs. After seeding, the cells were allowed to fully attach and the medium only changed the second day after isolation and cell seeding.

For RNA isolation, the isolated primary ECs were not resuspended in culture medium but snap-frozen in liquid nitrogen before undergoing the RNA isolation procedure.

### 2.4. Primary endothelial cell culture

The day after endothelial cell isolation, if ECs fully attached, the medium was gently changed to cell culture medium. If the cells did not fully attach yet, this procedure was postponed to the following day. Once grown to confluency on a 12-well, cells can be further passed onto one 6-well (P1), three 6-cm dishes (P2) and 10-cm dishes /T25 flasks (P3). The cells can be kept up to passage P4, requiring medium changes every 2-3 days. For BAT-ECs, the cells can typically be reseeded every seven days and about every four days for WATi-ECs.

### 2.5. muMEC cell culture

CI-muMEC (InScreenEx, INS-Cl-1004) were cultured according to manufacturer’s instructions on 2% gelatin-coated flasks in muMEC Medium (basal medium plus supplements, InScreenEx, INS-ME-1004) at 37°C 5% CO_2_. Medium was changed every 2-3 days and cells were subcultivated at 70-90% confluence.

### 2.6. Immunohistochemistry

Endothelial cells were seeded onto glass bottom dishes (ibidi, 81218-200) and allowed to reach 80-90% confluency before fixation. The cells were washed with 1X PBS, fixated with 4% paraformaldehyde (PFA) for 15 min at room temperature. After fixation, the cells were washed thrice with 1X PBS and stored in PBS or stained directly. For staining, the cells were blocked by incubating with blocking buffer (3% BSA, 2% FCS, 0.3% Tween®-20 in PBS) for 1 h at room temperature.

For immunohistochemistry from whole tissue, interscapular BAT was dissected and fixated overnight at 4°C in 10% formalin. Tissues were prepared for paraffin embedding using the automated embedding automate Microm STP-120 (Thermo Fisher Scientific). Samples were dehydrated in increasing isopropanol concentrations (70% for 3 h, two times 80% for 1 h each, two times 90% for 1 h each, 96% for 2h and two times and 100% for 2 h each) followed by two steps in xylene (two times 1 h each) to clear out the isopropanol. Samples were then incubated in molten paraffin grade 3 wax (two times 1 h each). The paraffin-infiltrated samples were embedded and casted into molds with liquid paraffin using the Microm EC 350 (Thermo Fisher Scientific) tissue embedding center. The tissues were sectioned into 7 µm slices using a Microm HM355 S microtome (Thermo Fisher Scientific) and mounted on Superfrost^TM^ Plus Adhesion microscope slides (New Erie Scientific LLC). The paraffin-embedded sections were deparaffinized by incubation in xylene (three times, 15 min each) and a 1:1 ratio of xylene and ethanol for 5 min. The samples were rehydrated in a graded ethanol series (three times 100% 2 min each, two times 90% 2 min each, two times 70 % 2 min each) followed by two water rinses for 2 min each. The samples underwent target antigen retrieval using citrate buffer (Sigma-Aldrich, C9999) for 10 min at 95°C in a water bath and allowed to cool down to room temperature (30-45 min). Slides were then washed with 1X PBS (twice for 2 min each, once for 10 min) and rinsed twice for 10 min (rinsing solution 0.2% gelatin plus 0.25% Triton X-100 in 0.5X PBS in H_2_O). Sample were encircled with hydrophobic pen and blocked in blocking buffer (3% BSA, 2% FCS, 0.3% Tween®-20, 0.25% Triton X-100 in PBS) overnight in a humidified chamber at 4°C. Subsequent staining steps for tissue sections were also performed in a humidified chamber.

Primary antibodies were diluted in blocking buffer with surface markers at 1:200 and ERG at 1:500 dilution. Primary antibodies were incubated in blocking buffer at 4°C overnight. Following washes with PBS, samples were incubated with Alexa-conjugated secondary antibodies at 1:500 dilution (Topro 1:2000) for 2h at 4°C. After washes with PBS, tissue sections were stained with DAPI (1:1000 in PBS) 10 min at 4°C. All samples were mounted using Fluoromount G (SouthernBiotech, 0100-01).

Representative images of were acquired using a Leica SP8 confocal microscope (Leica). For image processing, Fiji/ImageJ and Illustrator (Adobe) software were used.

### 2.7. Immunoblotting

For protein isolation, samples were lysed in RIPA buffer (Sigma, #R0278; 150 mM NaCl, 1.0% (v/v) IGEPAL CA-630, 0.5% sodium deoxycholate, 0.1% SDS, and 50 mM Tris, pH 8.0) supplemented with 1X EDTA-Free Complete Protease Inhibitor Cocktail (Roche, 11836170001), PhosSTOP™, Phosphatase Inhibitor cocktail mix (Roche, 4906845001) and 1 mM phenylmethylsulfonyl fluoride (Sigma-Aldrich, 10837091001). Cultured endothelial cells and muMECs were grown to about 90% confluency, washed with PBS and snap-frozen on dry ice. For protein isolation, 150 µL of lysis buffer were added and the cells detached and disintegrated by scraping down and transferred into a 1.5 mL tube.

For protein analysis from tissue, interscapular BAT, inguinal WAT or gonadal WAT tissue pieces were snap-frozen in liquid nitrogen directly after the harvest and stored at −80°C. Samples were manually mashed using Rotilabor®-micro pestle (Carl Roth, YE14.1) and the tissue integrity further disrupted by re-freezing on dry ice and grinding the tissue at least three times. Depending on the tissue type, the tissue mash was resuspended in different amounts of supplemented RIPA buffer (BAT and WATi: 600 µL, WATg: 300-400 µL buffer).

Protein lysates were stored on ice for 30 min and the samples were centrifuged for 15 min 17000 xg at 4°C. The supernatant was transferred into a new 1.5 mL tube and samples stored at −80°C. For tissue protein samples, the centrifugation and supernatant transfer step was performed twice. The protein concentration was determined using a BSA protein standard (Sigma-Aldrich, P0834-10X1ML) and ROTI Quant Bradford solution (Carl Roth, K015.1). The protein concentration at 595 nM was measured using a Spark® Multimode microplate reader (Tecan).

Protein samples were diluted in 4x Laemmli buffer (Thermo Fisher Scientific, J60015), boiled at 95°C for 10 min and stored at −20°C. The proteins were separated by SDS-PAGE (Tris-glycine gels with Tris/glycine/SDS buffer, Bio-Rad, 1610772) using 12% acrylamide gels (TGX FastCast Acrylamide Kit 12%, Bio-Rad, 1610175) according to manufacturers’ instructions. Samples and Precision Plus Protein™ Standards Dual Color (Bio-Rad, 161-0374) were loaded on 12% acrylamide gels and separated in an electric field of 90 V for varying time spans. Proteins were transferred onto nitrocellulose membranes (Trans-blot turbo RTA transfer kit, Bio-Rad, 1704271) using the Trans-Blot® Turbo™ Transfer System (Bio-Rad). Membranes were stained with Ponceau S solution (0.1% Ponceau, 5% acetic acid) to evaluate equal gel loading and transfer. Membranes were rinsed twice in TBS-T (0.01% Tween-20 in TBS) and blocked for 1 h in 5% milk in TBS-T. Membranes were incubated with primary antibodies at a 1:1000 dilution in 5% milk in TBS-T or 5% BSA in TBST overnight at 4°C. After washes and a consecutive 30 min 5% milk block, horseradish peroxidase-coupled secondary antibodies were incubated for 1.5 h at room temperature. To detect chemiluminescent protein signal, membranes were incubated in an ECL detection kit (Clarity™ Western ECL Substrate, Bio-Rad, 170-5061) and imaged using ChemiDoc™ MP Imaging System (Bio-Rad). Band intensities were quantified using the Image Lab software (Bio-Rad) and visualized using Photoshop (Adobe) and Illustrator (Adobe).

### 2.8. Gene expression analysis

RNA isolation from primary endothelial cells snap-frozen after the isolation was performed using RNeasy® Plus Micro Kit (Quiagen, 74034). RNA Isolation Kit was used according to the manufacturer’s instructions, incuding β-mercaptoethanol addition in the lysis step and eluting in 20 µL H_2_O. For primary ECs, cDNA synthesis was performed using High Capacity cDNA Reverse Transcription Kit (Thermo Fisher Scientific, 4368814).

RNA isolation from cultured endothelial cells and muMECs was performed using the NucleoSpin® RNA Plus Isolation Kit (Machery-Nagel, 74084.250). Cells were grown to about 90% confluency, washed with PBS and snap-frozen on dry ice. For RNA isolation, 350 µL of lysis buffer were added and the cells detached and disintegrated by scraping down. The cell lysate was collected in a 1,5 mL tube and the RNA Isolation Kit was used according to the manufacturer’s instructions, with one modification. During the elution step, the volume of elution buffer was reduced to 20 µL, which was subsequently reused for a second elution by adding it again to the RNA-binding column. RNA concentration was determined using a NanoDrop One, typically yielding concentrations between 50 and 100 ng/µL. For cDNA synthesis, 500 ng of RNA was used. cDNA synthesis was performed using the LunaScript RT Supermix Kit (New England Biolabs, E3010G).

For quantitative PCR (qPCR), TaqMan probes were employed to detect the target genes of interest. Beta-actin was selected as the reference gene due to its stable expression across various tissues and experimental conditions. The relative expression of the target genes was calculated using the ΔΔCt method, where the Ct value of the target gene was normalized to the Ct value of beta-actin to account for variations in RNA input or reverse transcription efficiency. Ct values were measured using QuantStudio 3 (Thermo Fisher Scientific).

### 2.9. Bulk RNA sequencing

For RNA-Seq, RNA was isolated from primary adult ECs from BAT, WATi and WATg depots as well as neonate primary and cultured (passage P1) BAT-ECs using the RNeasy® Plus Micro Kit (Quiagen, 74034). RNA Isolation Kit was used according to the manufacturer’s instructions, incuding β-mercaptoethanol addition in the lysis step. Bulk RNA sequencing way performed with Recipient Biomarker Technologies (BMK) GmbH. Briefly, RNA sample purity, concentration and integrity as well as library preparation quality were analysed by NanoDrop, Qubit 2.0 and Agilent 2100. The qualified library was sequenced using eukaryotic mRNA sequencing with a PE150 mode. Clean data of high quality was retrieved by filtering Raw data, thereby removing adapter sequence and low-quality reads. The obtained clean data was mapped to *Mus musculus* reference genome (Ensembl, Version GRCm38_release79) to generate mapped data. As library quality control, insert length and sequencing randomness were assessed on mapped data. RNA sequencing data set was analysed for gene expression quantification and principal component analyses (PCA) were performed on FPKM of each sample with similarity among them illustrated by reducing dimensionality into two or three principal components. Gene expression from adult BAT-ECs and WAT-ECs as well as neonate primary and cultured BAT-ECs was compared and differentially expressed genes (DEG) were determined using DESeq2 [18]. DEG were characterized by Fold Changes (FC) ≥2 and p-values <0.05. Gene Set Enrichment Analysis (GSEA) was performed.

### 2.10. Single-cell RNA-sequencing reanalysis

2.10.1. **Data description**

A published scRNA-seq dataset of cells isolated from the stromal vascular fraction of BAT (BAT-SVF) from mice housed at either thermoneutral (TN: 30 °C for 1 week), room temperature (RT: 22° C) or cold (5 °C for 2 days or 7 days) was downloaded from NCBI GEO database under accession number GSE160585, subsequently preprocessed and analyzed with Seurat v5 as detailed below.

#### 2.10.2. Preprocessing

Raw count matrices from GSE160585 were imported using the Read10X function in Seurat. Metadata provided with the dataset, as well as cluster identity information from supplementary tables, was incorporated into the Seurat object. For each sample, normalization was performed with SCTransform while regressing out cell cycle differences. The datasets were then integrated by selecting 3,000 features, computing anchors, and applying Seurat’s SCT-based integration workflow. Principal component analysis (PCA) was performed, followed by dimensionality reduction with UMAP based on the first 50 principal components. The number of components was determined by considering an inspection of the elbow plot, automated detection and expert review. A shared nearest-neighbor graph was constructed, and clustering was carried out at multiple resolutions (0.1–1.8).

Cluster annotation was guided by known marker genes, supplementary information from the original publication, and the identification of cluster-specific markers using FindAllMarkers. Cell identities were reassigned accordingly (e.g., endothelial cell subtypes, vascular smooth muscle cells, adipocytes, Schwann cells, pericytes, adipocyte progenitors, and immune cell subsets such as B cells, T cells, NK cells, monocytes, and macrophages). Custom color palettes were defined for visualization in UMAP plots. To focus on stromal and vascular populations, immune cells were collapsed into a single category and subsequently excluded, generating a subset object devoid of immune cells. The processed objects were saved as .rds files for downstream analyses.

#### 2.10.3. DE-Analysis

Analysis was conducted with the pre-integrated Seurat object provided as an .rds file from the preprocessing steps, which had already undergone quality control, normalization, and integration using the standard workflow in Seurat. We applied 50 dimensions throughout all analysis to ensure consistency across visualizations. Condition-specific embeddings were generated with the DimPlot function, and a custom color palette was applied to distinguish temperature treatments. For gene-level exploration, expression of set of genes was projected onto the UMAP embeddings using FeaturePlot, stratified by sample condition (temperature), and arranged into two-column grids with wrap_plots to enable direct comparison of expression patterns across conditions. Additionally lists for differentially expressed genes was created for pairwise comparisons of control and perturbations and provided as Excel files reporting p-value and log2FC of normalized RNA expression values from the corresponding assay, accompanying condition-specific Violin plots have been generated. DE testing used the default Wilcoxon rank-sum test with Bonferroni correction. Cell identities were confirmed with marker genes. Violin plots of gene expression has been reported for different gene groups in addition.

The preprocessing and analysis scripts have been deposited the following Github repository: https://github.com/stephanmg/shamsi-BAT-re-analysis

### 2.11. Quantification and statistical analysis

Statistical analysis was carried out using PRISM 10 (GraphPad Software) on the raw data. Comparisons between two groups were made using a t-test, unless otherwise specified in the figure legends. A p-value of p<0.05 was considered statistically significant. Data are presented as dot plots, with vertical bars representing the means, unless otherwise noted in the figure legends.

## 3. Results

### 3.1. Isolation of endothelial cells (ECs) from neonatal and adult adipose tissue

This protocol was developed to isolate ECs from interscapular brown adipose tissue (BAT), inguinal white adipose tissue (WATi) and gonadal white adipose tissue (WATg). A step-by-step procedure of the isolation process is presented in Figure 1A.

**Figure 1:**
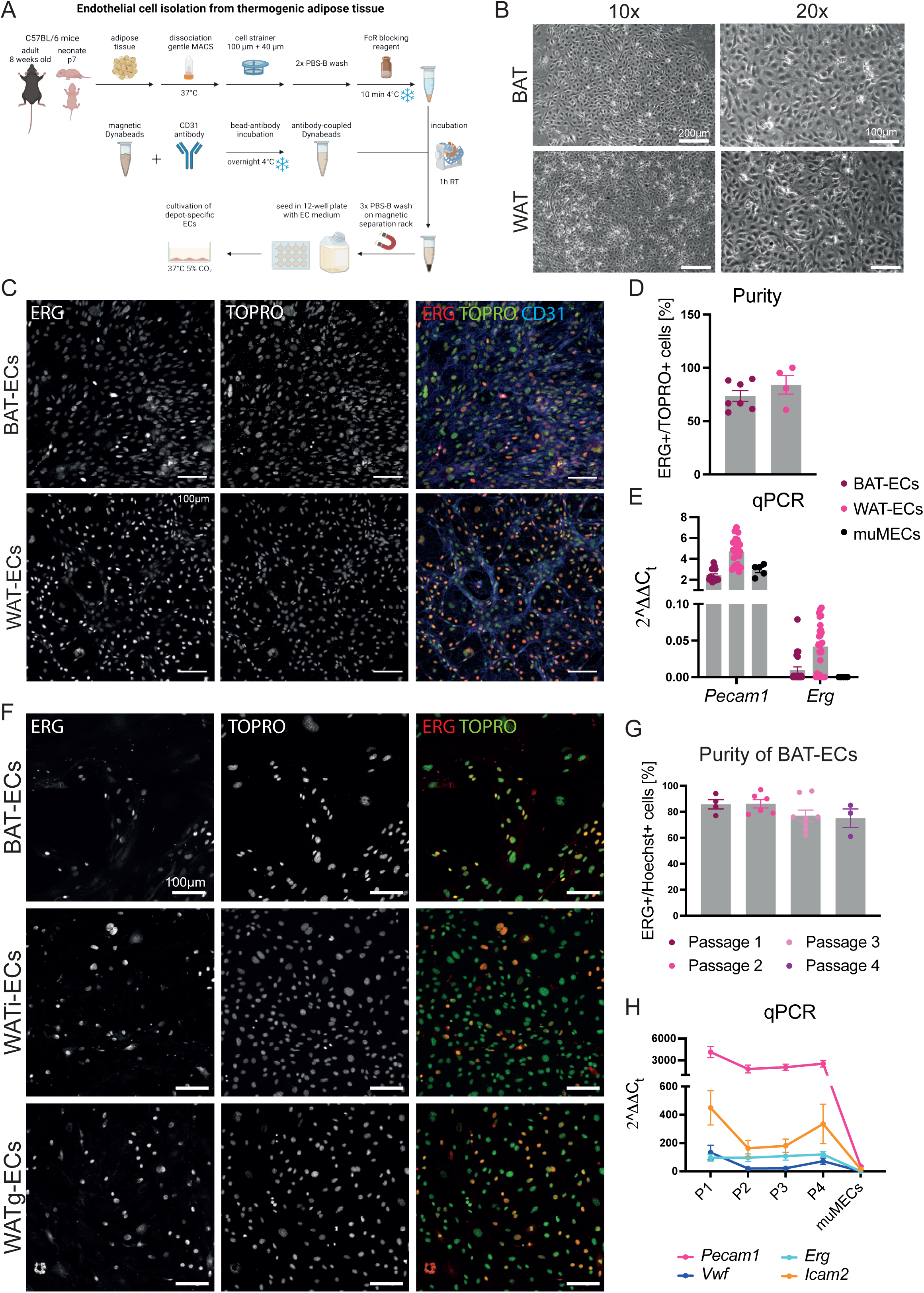
Our endothelial isolation protocol gives rise to high purity of adipose ECs. **A** Schematic representation of the major steps of endothelial cell isolation from thermogenic adipose tissue. **B** Brightfield images of endothelial cells isolated from brown adipose tissue (top panels) and white adipose tissue (bottom panels) at different magnifications (10X, 20X). **C** Confocal images of BAT-ECs (top) and WAT-ECs (bottom) isolated from neonatal mice and labelled for ERG, TOPRO and CD31 (Pecam1). These images display the high purity of cultures. **D** Purity of neonatal endothelial cultures (passage 1) of BAT (n=7 independent samples) and WAT (n=4 independent samples) determined from culture dishes labelled for ERG and TOPRO. **E** mRNA levels of the endothelial markers *Pecam1* and *Erg* at early passages of BAT-ECs (n=23 independent samples) and WAT-ECs (n=28 independent samples) in comparison to the immortalized murine EC line, muMECs (n=5 independent samples). This displays high endothelial marker expression of BAT-ECs and WAT-ECs. **F** Representative confocal Images of BAT-ECs, WATi-ECs and WATg-ECs isolated from adult mice and stained for ERG and TOPRO, displaying high purity of the endothelial cultures. **G** Quantification of BAT-EC purity at different passages (n≥3 independent samples), showing that purity only mildly drops from one passage to the other. **H** qPCR results of endothelial markers (*Pecam1, Erg, Vwf, Icam2*) of BAT-ECs at different passages (n≥3 independent samples), representing high levels of endothelial markers at different passages in comparison to muMECs.

In brief, AT was initially dissociated by chopping and enzymatic digestion. The resulting cell suspension was then filtered through 100 µm and 40 µm cell strainers to obtain a single-cell suspension. After washing, nonspecific binding was blocked. Subsequently, the single-cell suspension was incubated with CD31-coupled magnetic beads. Using magnetic separation racks, the bead-labeled cells were washed and then resuspended in EC culture medium (DMEM-F12 supplemented with 20% FCS, endothelial growth factors, and antibiotics).

The isolated ECs were seeded onto gelatin-coated 12-well plates at a density of approximately 200,000 cells per well. For the isolation of ECs from adult fat depots, the tissues do not need to be pooled, allowing for the comparison of ECs from mice of different genotypes. In contrast, for the isolation of ECs from neonatal adipose tissue, fat depots from 3–5 neonates were pooled to ensure a sufficient number of ECs for culture. For neonatal isolations, the protocol yielded an average of 305 000 (± 95 000) BAT-ECs and 470 000 (± 260 000) WATi-ECs, when fat depots of four pups were pooled. In contrast, EC isolations from adult mice resulted in an average yield of 120 000 (± 90 000) BAT-ECs, 1 530 000 (± 350 000) WATi-ECs and 1 430 000 (± 470 000) WATg-ECs per single fat depot. After seeding, the cells were allowed to fully attach for two days before the culture medium was changed. At this stage, the cells already exhibited the characteristic cobblestone morphology of ECs (Supplementary Figure 1A).

Once the cells reached confluency in a 12-well plate, they were passaged to a 6-well plate (passage 1). Upon reaching confluency again, the cells were split into three 6-cm dishes (passage 2). In the third passage, cells from one confluent 6-cm dish were seeded onto a 10-cm dish or T25 flask. The cells could be maintained up to passage P4, with medium changes required every 2–3 days. BAT-ECs could be reseeded approximately every seven days, whereas WAT-ECs reached confluency in about four days.

### 3.2. The protocol gives rise to a high purity of ECs

A fast and cost-effective quality control method for EC cultures is the assessment of cell morphology. ECs are typically characterized by a cobblestone-like shape, which serves as a criterion for high-quality EC cultures. For the successful culture of ECs, it is essential to minimize contamination by mesenchymal cells, particularly fibroblasts. Fibroblasts exhibit a lower doubling rate than ECs and can rapidly overgrow EC cultures within a single passage (Supplementary Figure 1B). In contrast to ECs, fibroblasts look spikier and do not grow in monolayers. The purified BAT- and WAT-ECs isolated using our protocol display a uniform cobblestone monolayer (Figure 1B), suggesting a high degree of endothelial purity.

While flow cytometry is commonly used to assess EC purity in other protocols [19,20], this approach was not suitable in our case, as the magnetic beads used for EC isolation are not compatible with consecutive flow cytometry. Therefore, we implemented an alternative method to evaluate and quantify cell purity. We seeded approximately 40,000 isolated cells from the first passage onto small staining dishes and subsequently performed immunofluorescent staining. ECs were identified using an antibody specific for the ETS transcription factor ERG, which labels the cell nucleus [21]. By co-staining with a nuclear marker, such as TOPRO or Hoechst, we were able to distinguish and quantify double-positive cells as ECs (Figure 1C). Using this approach, our isolation protocol consistently achieved a purity of approximately 75% for brown adipose tissue-derived ECs (BAT-ECs) and 85% for ECs isolated from inguinal white adipose tissue (WAT-ECs) in the first passage (Figure 1D).

In addition to immunofluorescence-based purity assessment, we isolated RNA from the cultured cells to examine the expression of key blood endothelial markers, including *Pecam1* and *Erg.* We compared these primary ECs to early-passage commercially available, immortalized murine capillary ECs isolated from the mouse lung (muMECs, InScreenEx) (Figure 1E): BAT-ECs exhibited slightly decreased expression of *Pecam1*, but clearly increased *Erg*, while WAT-ECs showed more abundant transcript levels for both. Additionally, BAT- and WAT-ECs displayed higher endothelial marker expression compared to muMECs, further underlining the endothelial identity of BAT- and WAT-ECs (Supplementary Figure 1C).

In summary, the EC isolation protocol described here, yields highly pure endothelial cells. Further, the findings confirm that our protocol not only enables the isolation of ECs from AT but also maintains endothelial characteristics under our culture conditions, providing an additional quality criterion for this method.

### 3.3. The isolation protocol is effective in adult mice

Consistent with the results obtained from neonatal fat depot isolations, the protocol achieves high EC purity in adult tissue-derived ECs. Staining for the endothelial-specific nuclear marker ERG and comparing it to total nuclear staining with TOPRO or Hoechst revealed a BAT-EC purity of approximately 82%, which declined slightly from passage 1 to passage 4 (Figure 1F, G).

To further validate the endothelial identity of the isolated cells, we quantified transcript levels of canonical EC markers (*Pecam1*, *Erg*, *Icam2*, *Vwf*) in BAT-ECs and compared them to those in the immortalized murine EC line (muMECs). Further, we followed the marker expression over four passages (Figure 1H). All canonical endothelial markers were highly expressed in primary BAT-ECs, whereas their expression levels in muMECs were significantly lower or barely detectable.

These findings confirm the robustness and reliability of our protocol for isolating ECs from adult adipose tissue and further emphasize the advantages of using primary ECs over immortalized endothelial cell lines for physiologically relevant studies.

### 3.4. ECs isolated from BAT express depot-specific signatures

Next, we assessed the depot-specific transcriptomes of ECs. Using our protocol, we isolated ECs from adult BAT, WATi, and WATg. RNA was extracted immediately from the primary cells and subjected to RNA sequencing. Principal component analysis (PCA) showed that overall gene expression was highly similar among ECs isolated from the same fat depot (Figure 2A). Along principal component 2 (PC2), WATi-ECs, originating from a beige-fat depot, clustered between BAT-ECs, representing pure brown fat, and WATg-ECs, representing the whitest depot. Along PC1, however, WATi-ECs clustered far from both BAT- and WATg-ECs. When examining canonical endothelial markers, WATi-ECs displayed very low expression levels, and were therefore excluded from further analysis (Figure 2B). By contrast, BAT-ECs and WATg-ECs expressed high levels of canonical endothelial markers along with depot-specific markers *Gpihbp1* and *Plvap*, respectively (Figure 2B). Subsequent comparative analysis focused on BAT-ECs versus WATg-ECs. KEGG pathway analysis confirmed the specificity of our approach, with significantly enriched pathways related to adipocyte metabolism, including PPAR signaling, insulin signaling, thermogenesis, fatty acid metabolism, and regulation of lipolysis (Figure 2C). These results suggest that BAT-ECs may directly support adipocyte metabolic function.

**Figure 2:**
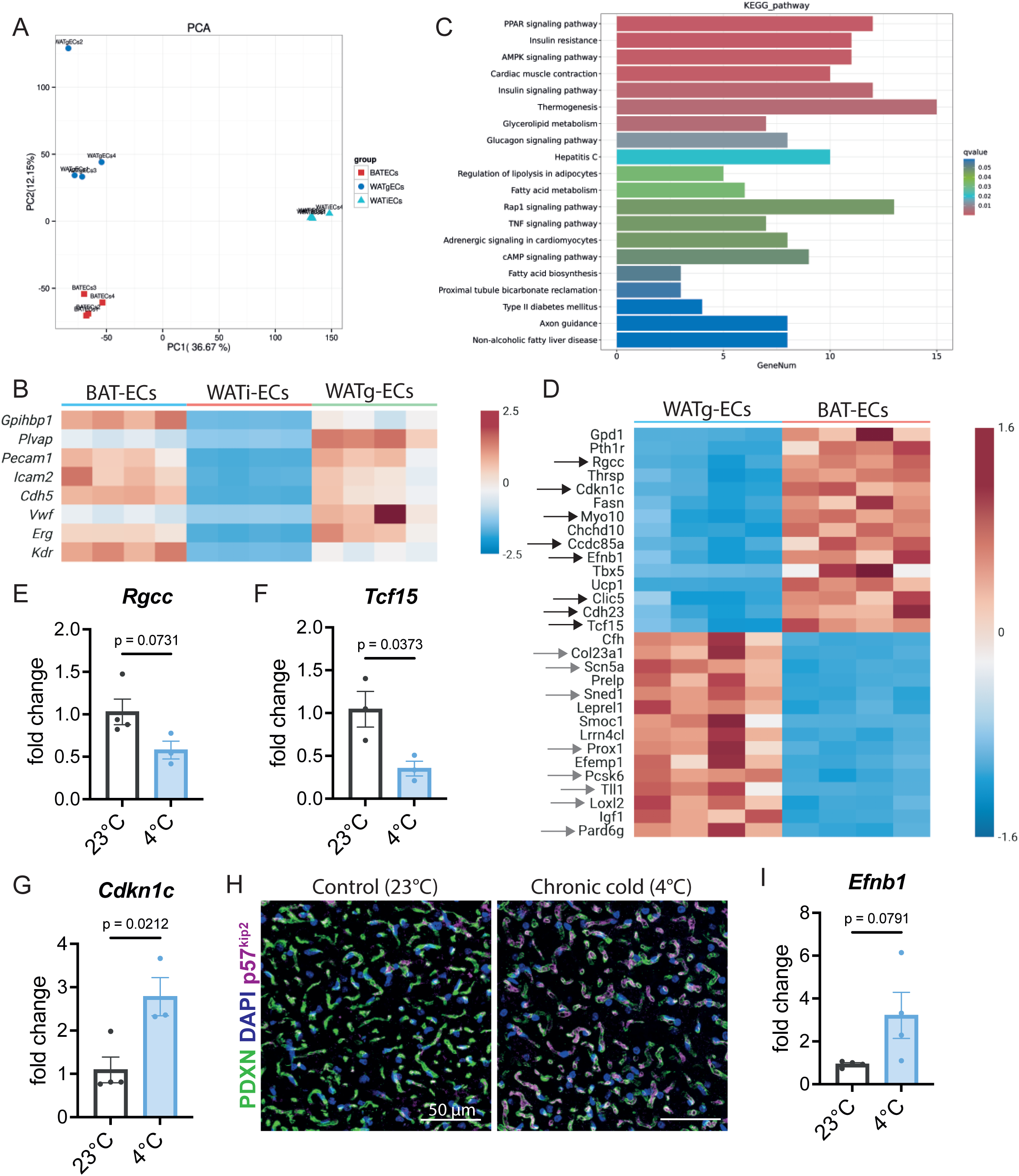
Transcriptomic analysis of isolated BAT-ECs defines novel BAT-specific EC markers. **A** Principal Component Analysis (PCA) comparing the endothelial transcriptome of 4 replicates of each fat depot (BATECs, WATiECs, WATgECs) representing low variability on y-axis (PC2) but high deviation of WATiECs on x-axis (PC1). **B** Heatmap of endothelial marker gene expression measured through mRNA sequencing of BAT-ECs, WATi-ECs and WATg-ECs, showing that WATi-ECs express endothelial markers only at very low levels. BAT-EC marker *Gpihbp1* is highly abundant in BAT-ECs, while WAT-EC endothelial marker *Plvap* is strongly transcribed in WATg-ECs. **C** KEGG pathway analysis using all differentially upregulated genes in BAT-ECs compared to WATg-ECs. **D** Heatmap of the 15 most significantly enriched genes in BAT-ECs compared to WATg-ECs and the 15 most significantly upregulated genes in WATg-ECs compared to BAT-ECs. **E** mRNA levels of *Rgcc*, **F** *Tcf15* and **G** *Cdkn1c* in BAT-ECs of C57Bl6 mice housed at 23°C (n≥3 independent samples) and 4°C (n=3 independent samples) detected using qPCR. **H** Podocalyxin (PDXN), DAPI and p57^kip2^ immunofluorescence labeled BAT of C57Bl6 mice housed at 23°C or 4°C, presenting increased and endothelial specific p57^kip2^ fluorescence in the 4°C condition. **I** Transcript levels of *Efnb1* in BAT-ECs of C57Bl6 mice housed at 23°C or 4°C (n=4 independent samples each condition) detected using qPCR. For **E**, **F**, and **G** data represent mean ± s.e.m.; two-tailed unpaired t-test. The numerical data and *P* values are provided in the figure. *P* values lower that 0.05 are considered as significant.

To further characterize depot-specific features, we identified the 15 most highly enriched genes in BAT-ECs compared to WATg-ECs (Figure 2D). To confirm endothelial specificity, we analyzed published single-cell RNA-seq data [22]. This revealed that eight of the top 15 genes were either enriched (*Rgcc, Tcf15*) or exceptionally expressed (*Cdkn1c, Ccdc85a, Myo10, Efnb1, Clic5, Cdh23*) in BAT-ECs (Supplementary Figure 1D-S). To prioritize functional candidates, we then assessed whether these genes were regulated upon BAT activation by chronic cold exposure (7 days at 5°C) [22] (Supplementary Figure 3). Indeed, *Rgcc*, *Tcf15*, and *Cdkn1c* were significantly regulated under cold exposure (Supplementary Figure 2). We therefore focused on these genes and evaluated whether they have been previously linked to endothelial or adipose tissue biology.

Rgcc (Regulator of Cell Cycle; also known as RGC-32) is broadly expressed and has been reported to enhance proliferation by repressing cell-cycle inhibitors, but in some cancer cells it inhibits mitosis and blocks tumor growth. In ECs, RGC-32 has been described as a capillary marker and as a hypoxia-induced regulator that limits proliferation, migration, and angiogenesis [6,23] . Intriguingly, RGC-32 knockout mice display increased browning and energy expenditure, protecting against diet-induced obesity [24]. In our reanalysis, *Rgcc* expression decreased significantly under cold exposure (Supplementary figure 2A). We confirmed this by measuring *Rgcc* in freshly isolated BAT-ECs before and after cold exposure (Figure 2E). However, analysis of whole AT lysates show that RGC-32 accumulates under cold exposure in BAT, WATi and WATg (Figure 3A-F).

**Figure 3:**
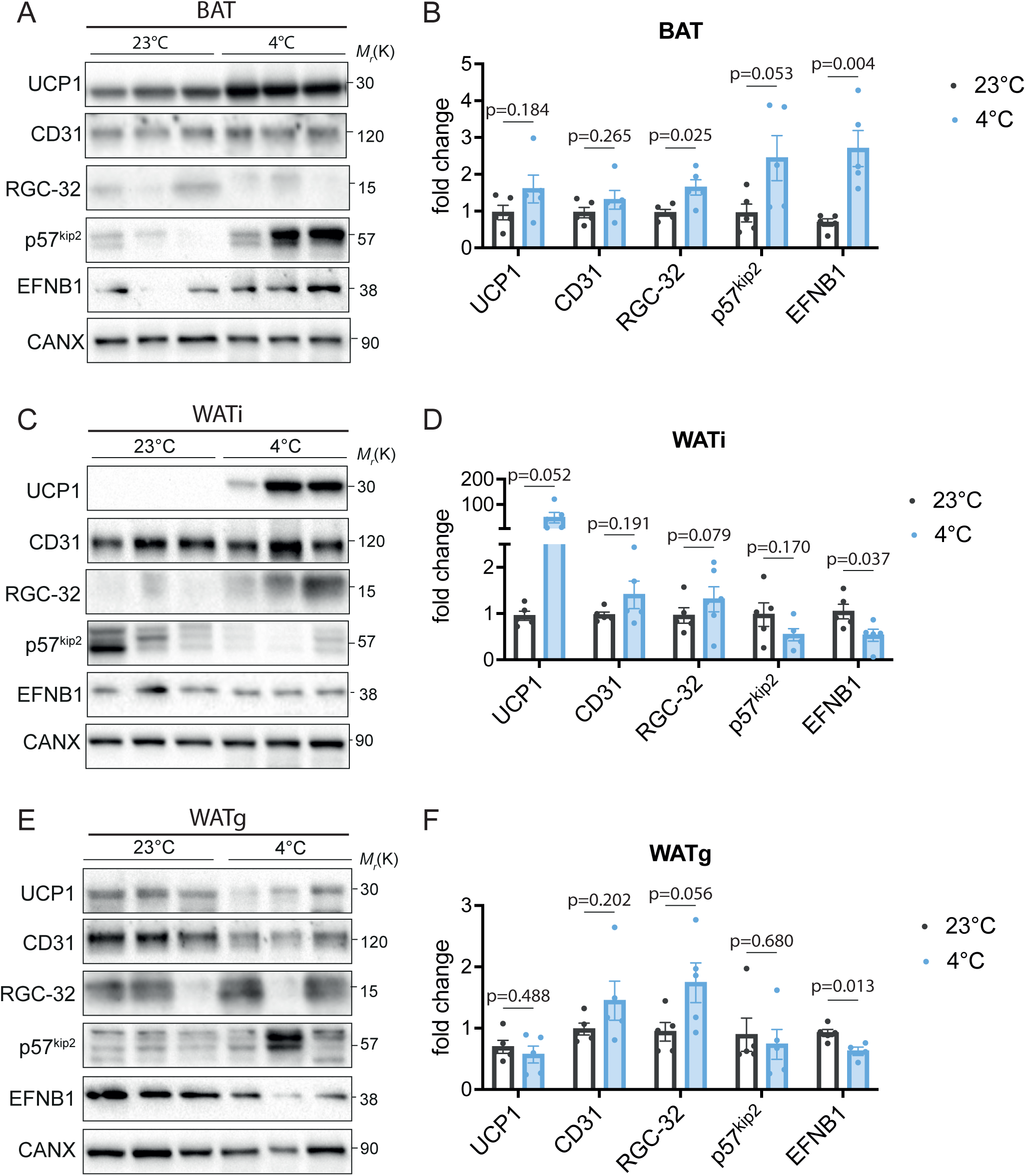
Our newly defined BAT-EC markers are regulated by cold exposure. Immunoblots representing RGC-32, p57^kip2^ and EFNB1 protein levels at 23°C and 4°C in BAT (**A**), WATi (**C**) and WATg (**E**). Further the blot displays CD31 to assess endothelial content, UCP1 to validate the response to 4°C and Calnexin (CAXN) as housekeeping protein. Quantification of protein levels of UCP1, CD31, RGC-32, p57^kip2^ and EFNB1 in BAT (**B**), WATi (**D**) and WATg (**F**) at 23°C (n=5 independent samples) and 4°C (n=5 independent samples). For **B**, **D** and **F** data represent mean ± s.e.m.; two-tailed unpaired t-test. The numerical data and *P* values are provided in the figure. *P* values lower that 0.05 are considered as significant.

TCF15 (Transcription Factor 15) regulates fatty acid uptake in cardiac ECs, where it heterodimerizes with MEOX2 to control cardiomyocyte metabolism [25]. Coppiello et al. showed that TCF15 and MEOX2 are highly expressed in ECs of tissues with strong reliance on fatty acid metabolism, including BAT, WAT, and heart [25]. Consistent with this, our bulk RNA-seq data detected abundant *Meox2* in both BAT- and WATg-ECs, although not differentially regulated. Surprisingly, reanalysis of single-cell RNA-seq revealed significant downregulation of *Tcf15,* but not *Meox2* upon cold exposure, which we confirmed in isolated BAT-ECs (Figure 2F). Together, these results suggest that TCF15 and MEOX2 serve as adipose-specific EC markers, that are downregulated during BAT activation.

Cdkn1c (Cyclin-dependent kinase inhibitor 1c; p57^Kip2^) is a well-established negative regulator of cell cycle progression. Previous studies demonstrated its high abundance in BAT compared to WATi and WATg [26]. In our analysis, *Cdkn1c* was specifically enriched in BAT-ECs and upregulated after we exposed mice to 4°C, prompting further validation (Figure 2G). Immunostaining of BAT from mice housed at room temperature or after one week of cold exposure revealed strong p57^kip2^ signal in cold-exposed BAT, colocalizing with the endothelial marker Podocalyxin (PDXN) (Figure 2H). Additionally, immunoblotting of whole AT lysates showed increased p57^kip2^ levels in BAT upon cold exposure, while it was barely changed in WATi and WATg (Figure 3A-F). These findings suggest that p57^kip2^ acts as an endothelial-specific regulator in thermogenic adipose tissue, potentially contributing to the endothelial response to cold-induced thermogenesis.

Although *Efnb1* was not significantly regulated by cold exposure in published snRNA-seq data [9], we explored it further as Ephrins have been shown to be important in ECs. Ephrin-B1 (Efnb1), a ligand of EphB receptors, regulates cell–cell communication and angiogenesis. Low *Efnb1* levels in murine AT have been linked to obesity and inflammation [27]. We found that EFNB1 protein abundance significantly increased in BAT after cold exposure, consistent with elevated expression in isolated BAT-ECs (Figure 2I and Figure 3A, B). Conversely, EFNB1 decreased in WATi and WATg under the same conditions (Figure 3B-F). This opposing regulation suggests depot-specific roles for EFNB1 in vascular adaptation, potentially supporting angiogenesis in thermogenically active BAT while being downregulated in WAT.

In summary, our approach identified several adipose-specific EC markers, including RGC-32, TCF15, MEOX2, p57^kip2^, and EFNB1. Many of these genes are dynamically regulated during BAT activation, highlighting specialized roles for ECs in supporting thermogenic adipose tissue.

### 3.5. Transcriptome of freshly isolated versus cultured ECs

The aim of this study was not only to validate our BAT-EC isolation method and identify novel BAT-EC-specific marker genes, but also to analyze potential changes during culture. Thus, we compared freshly isolated BAT-ECs with BAT-ECs cultured over one passage. PCA showed that freshly isolated BAT-ECs clustered tightly along PC1 and PC2, indicating a highly similar transcriptome, whereas cultured BAT-ECs clustered together on PC1 but shifted markedly along PC2, suggesting that culture conditions diversify their transcriptome (Figure 4A). KEGG analysis revealed significant upregulation of pathways related to focal adhesion and cell cycle regulation, which reflects the expected adaptation of ECs to *in vitro* conditions, where they must proliferate, grow, and adhere to culture dishes. Pathways such as PI3K–AKT and mTOR signaling, which support EC proliferation and growth, were also enriched, as well as metabolic pathways associated with nucleotide and amino acid synthesis— critical for cellular mass doubling (Figure 4B). Interestingly, thermogenesis-related genes were also upregulated, suggesting that cultured cells retain elements of depot-specific identity (Figure 4B).

**Figure 4:**
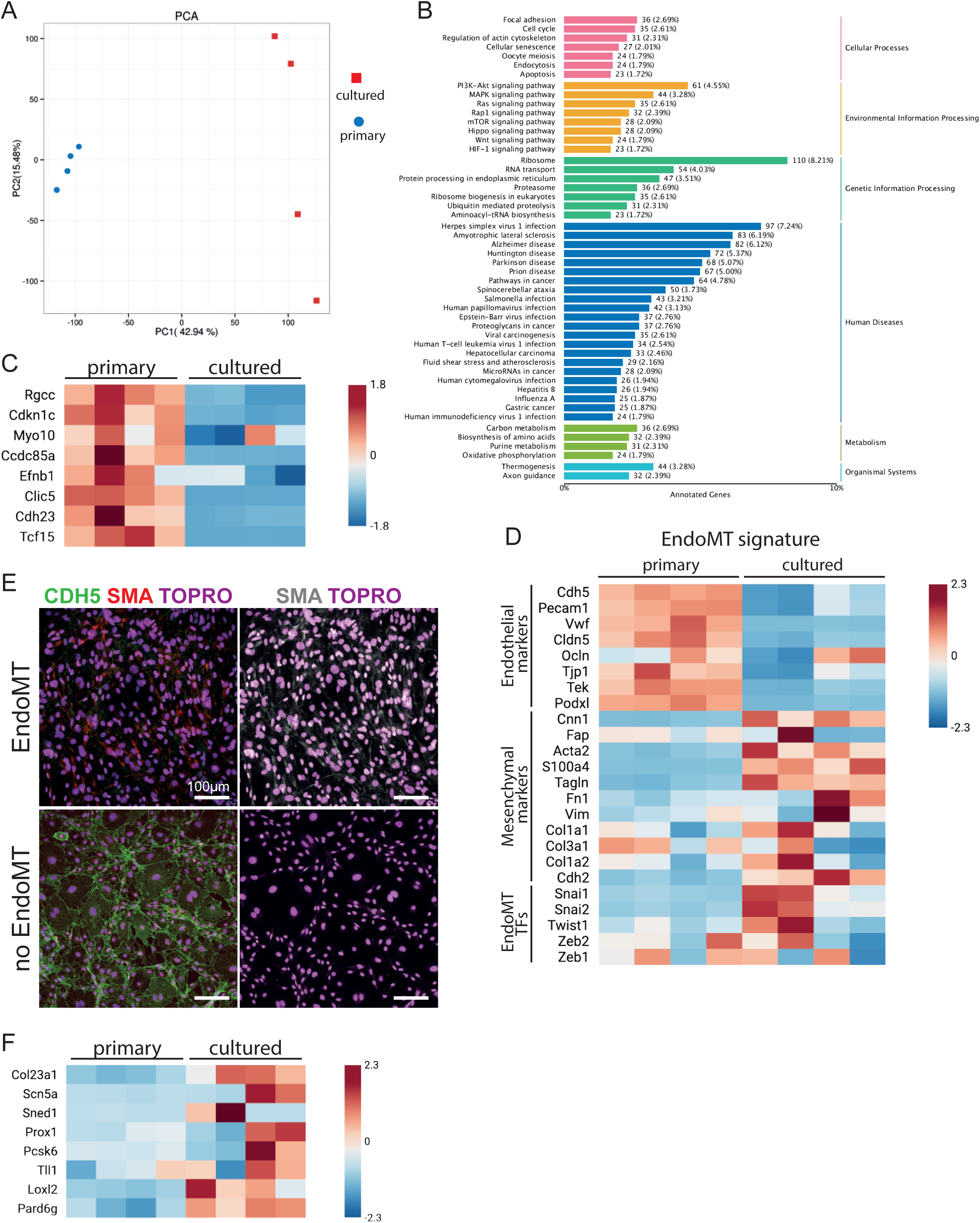
Cultured BAT-ECs lose their specific markers and gain WAT-EC characteristics. **A** Principal Component Analysis (PCA) of 4 replicates of cultured and primary endothelial cells isolated from BAT. This PCA shows that the transcriptome of primary BAT-ECs clusters closely together, while cultured BAT-ECs differ clearly in PC2 and separate from primary BAT-ECs on PC1. **B** KEGG pathway analysis using all significantly upregulated genes in BAT-ECs cultured for one passage compared to primary BAT-ECs. **C** Heatmap comparing the transcript levels of defined BAT-EC markers from primary BAT-ECs versus cultured BAT-ECs. **D** Heatmap of genes involved in endothelial to mesenchymal transition (EndoMT) revealing that endothelial markers decrease under culture conditions, while mesenchymal marker and EndoMT transcription factors increase. This suggests that BAT-ECs undergo EndoMT *in vitro*. **E** Representative confocal images of BAT-ECs culture undergoing EndoMT (upper panels) or with no signs of EndoMT (lower panels). EndoMT is identifies by the loss of CDH5 staining and gain of smooth muscle actin (SMA) staining. **F** Heatmap of mRNA levels of defined WAT-EC markers in primary BAT-ECs compared to cultured BAT-ECs. This confirm that BAT-ECs transition to a WAT-ECs transcriptome in culture.

To explore this further, we examined mRNA levels of the BAT-EC marker genes newly defined in this study. All were significantly downregulated (Figure 4C). Given that many of these markers suppress cell cycle progression, their downregulation is consistent with the proliferative drive of cells under culture conditions. Moreover, *in vitro* BAT-ECs lack blood flow, their native microenvironment, and surrounding smooth muscle cells, all factors that promote dedifferentiation and can trigger an endothelial-to-mesenchymal transition (EndoMT) [15]. Indeed, transcriptomic analysis indicated increased EndoMT signatures. This means endothelial marker genes are downregulated, while mesenchymal genes and transcriptions factors known to drive epithelial to mesenchymal transition (EMT) are upregulated (Figure 4D). We further determined EndoMT on protein levels and stained BAT-EC cultures with the endothelial marker VE-Cadherin (CDH5) and the mesenchymal protein α-smooth muscle actin (SMA). While early cultures (passage 1) mainly showed strong CDH5 staining and no SMA, late passages (passage 4 and later) barely displayed CDH5 fluorescent staining, but increased SMA levels (Figure 4E).

We next hypothesized that, as BAT-ECs lose BAT-specific markers in culture, they may acquire WAT-specific ones. To test this, we revisited our comparison of BAT-ECs and WATg-ECs and identified the 15 genes most enriched in WATg-ECs (Figure 2D). Analysis of published single-cell RNA-seq data [22] revealed that many of these genes were endothelial-enriched, particularly in lymphatic ECs (Supplementary Figure 3). For example, *Prox1*, a master transcription factor of lymphatic EC identity and widely used driver in lymphatic EC studies was among the most enriched [28] . When these WATg-EC markers were examined in our datasets, they were virtually absent in freshly isolated BAT-ECs but strongly upregulated in cultured BAT-ECs (Figure 4F).

Together, these findings demonstrate that BAT-ECs undergo profound transcriptomic remodeling in culture, characterized by loss of BAT-specific markers and acquisition of WATg- and lymphatic EC-like features. This suggests that BAT-ECs undergo a form of “whitening” under culture conditions, with a shift toward a lymphatic EC transcriptome. Moreover, these results raise the possibility that highly specialized BAT-ECs may derive from, or revert to, a WAT-EC state with lymphatic features under certain conditions.

## 4. Discussions

Studying the vasculature in adipose tissue faces major hurdles [29]. In particular, the study and identification of angiokines and batokines along with their mechanisms of action is extremely challenging. *In vivo* studies require genetically modified mice and sophisticated and expensive technology [30,31]. Additionally, the results are highly dependent on the endothelial-specific Cre-recombinase. As there is so far no recombinase that targets only the vasculature of AT, the interpretation of the results is extremely challenging and it is hard to exclude the contribution from other organs.

Thus, isolating and culturing ECs from AT enables in-depth investigation of their unique characteristics, signaling pathways, and interactions within the microenvironment of brown, beige and white fat. However, due to the small size and complex structure of adipose depots, isolating pure populations of viable ECs and bringing them into culture presents a distinct set of challenges.

While several published protocols describe EC isolation, none fully met the specific requirements. Many focus on isolating ECs from the lung, heart, or aorta, often involving complex perfusion techniques that are difficult to implement in small adipose depots [19,20,32–34]. Others describe EC isolation from human or mouse tissues but lack a focus on long-term culture conditions, a critical aspect for functional studies [35–37]. Additionally, commercially available primary or immortalized ECs predominantly originate from the mouse lung, human umbilical cord, or skin. These ECs lack the necessary characteristics of AT-derived ECs, making them unsuitable for investigating the signaling mechanisms between ECs and adipocytes.

Our protocol yields approximately 80% pure ECs from both BAT and WAT. Moreover, this study revealed several novel BAT-specific endothelial markers: *Rgcc*, *Cdkn1c*, *Tcf15*, *Meox2* and *Efnb1*.

Rgcc (or RGC-32) is a VEGF and hypoxia-induced protein that inhibits angiogenesis[23] . Interestingly, the RGC-32 KO mouse displays increased browning and energy expenditure, protecting the mouse from obesity [24]. The authors did not find a major functional change in RGC-32 knockdown adipocytes, suggesting that other cells in AT are affected by the loss of RGC-32. Our reanalysis of single-cell RNASeq data showed that endothelial *Rgcc* levels significantly drop in mice exposed to cold. Thus, deletion of RGC-32 in the endothelium might enable an otherwise restricted angiogenic response. Increased vascularization supports BAT [38] and together this could explain the maintained browning and resistance to HFD-induced obesity in the RGC-32 KO mice. On the other hand, we find increased protein levels of RGC-32 in BAT tissue upon cold exposure. As RGC-32 is induced by VEGF or hypoxia, which are high under cold exposure [39], it is thus not surprising that RGC-32 is upregulated. This apparent discrepancy between transcriptomic downregulation in ECs and protein upregulation at the tissue level highlights the complexity of RGC-32 regulation and warrants further investigation.

P57^kip2^ (Cdkn1c) is known to inhibit cell cycle progression. Mutations in this gene are common in the growth restriction disorder Silver Russell Syndrome, IMAGe Syndrome and Beckwith Wiedemann Syndrome [40]. Besides the strong growth retardation, these patients show significantly reduced adiposity. A mouse model of Cdkn1c overexpression, mimicking the described disorders, has substantially more BAT, while the Cdkn1c KO mouse fails to develop BAT [40]. In particular, the Cdkn1c KO mouse fails to accumulate PRDM16 and thus to develop brown/beige adipocytes. In human adipose-derived stem cells p57^Kip2^ was shown to inhibit key cell cycle molecules, thereby inducing quiescence or even senescence [41]. Another related cell cycle inhibitor, p27^Kip1^, which shares the same degradation pathway, has been shown to restrict proliferation in mature brown adipocytes and, consequently, affect thermogenic adaptation in BAT. This study also reported that p57^Kip2^ is exclusively expressed in BAT compared to gonadal white adipose tissue (WATg) and inguinal WAT (WATi) [26]. Despite this, only a single study has suggested that Cdkn1c may be enriched in endothelial cells (ECs). Specifically, p57^Kip2^ was described as a downstream target of KLF2 and KLF4, and its expression was upregulated by laminar flow, though this regulation was specific to lymphatic endothelial cells [42]. Notably, blood ECs did not show a similar flow-induced increase in Cdkn1c expression [42].

Ephrin-B1 (encoded by Efnb1) is a transmembrane ligand within the B-class ephrin family, engaging in bidirectional signaling with EphB receptors to orchestrate cell–cell communication. In ECs, ephrin–Eph interactions are pivotal in vascular development and angiogenesis: for instance, ephrin-B1 is expressed in arterial and venous endothelial cells and contributes to capillary sprouting and vessel patterning during vascular morphogenesis [43–45]. Specifically, human coronary artery ECs express ephrin-B1 protein, and its expression can be induced by inflammatory stimuli [27]. Beyond vascular biology, ephrin-B1 was also detected in AT. Expression of ephrin-B1 is notably reduced in adipose tissue—especially in mature white adipocytes—of obese mice, and its downregulation exacerbates adipose inflammation. Overexpressing ephrin-B1 in adipocytes, on the other hand, suppresses inflammatory responses, underscoring a protective role in AT homeostasis [46]. In our study, we observed a striking depot-specific regulation of Ephrin-B1 (EFNB1) in ECs. In BAT, where *Efnb1* in ECs is already high at baseline, cold exposure led to a further upregulation. By contrast, in WATi and WATg, where endothelial *Efnb1* is relatively low, cold exposure resulted in a significant downregulation. This opposing regulation suggests that EFNB1 may play a distinct role in the vascular adaptation of different adipose depots.

One possible explanation is that in BAT, endothelial EFNB1 upregulation supports angiogenesis and vascular remodeling in response to thermogenic activation, thereby facilitating the increased nutrient and oxygen demands in cold-activated BAT. In WAT, however, downregulation of EFNB1 is likely not mediated by ECs, but other cells, e.g. mature adipocytes [46]. This opposing regulation highlights the depot-specific signaling and suggests that Ephrin–Eph signaling contributes not only to vascular plasticity but also to the functional divergence between BAT and WAT vasculature.

We developed our protocol not only to isolate and enrich BAT-ECs, but also to culture and expand them. Here we made several interesting findings. ECs isolated from BAT lose our defined BAT-EC marker expression. Several of these markers prevent cell cycle progression. At the adult stage when we isolate BAT-ECs, ECs are mainly quiescent and do not actively proliferate. Thus, it makes sense that we found several cell cycle inhibitors. When we bring these cells in culture, the cells are forced to proliferate and have to undergo a rapid angiogenic switch. In culture, ECs partly lose their organ-specific expression pattern, as the transcriptome is highly influenced by the organ/microenvironment the cells reside in [6]. Further, the transcriptome of cultured BAT-ECs suggests that they undergo EndoMT, a process frequently reported as a consequence of *in vitro* conditions [15]. Rather than viewing this as a limitation, we propose that such remodeling represents a necessary step for ECs to exit their highly specialized *in vivo* state and enable robust proliferation *in vitro*. In this context, the acquisition of mesenchymal-like features through EndoMT [15,47,48] may be interpreted as part of a broader adaptive program that supports survival and expansion under culture conditions. Interestingly, in high passage BAT-EC cultures we observed cells containing several small lipid droplets, reminiscent of brown adipocytes. One possible interpretation is that ECs, after transitioning through mesenchymal-like state, may further acquire adipocyte-like features. In line with this, the EndoMT signature observed in cultured BAT-ECs could represent an intermediate step toward a more multipotent mesenchymal phenotype, from which adipocytes can also be derived [49]. While highly speculative, this raises the intriguing possibility that cultured BAT-ECs may retain unexpected lineage plasticity.

BAT-ECs in culture quickly adopt a WATg-EC transcriptome. In particular, we found increased expression of canonical lymphatic EC markers within cultured BAT-ECs. This may indicate that BAT-ECs, when forced to proliferate, leave their highly specialized state and adopt a more permissive transcriptional program resembling WAT-ECs. While speculative, it is noteworthy that endothelial cells can exhibit lineage plasticity, and lymphatic transdifferentiation has been described in certain contexts [50]. Thus, the lymphatic-like features of cultured BAT-ECs may reflect such a plastic response, though this possibility requires further investigation.

In summary, our work establishes a protocol for the isolation and culture of ECs from brown and white adipose depots, enabling functional and molecular studies of their unique properties. Using this approach, we identified several BAT-specific endothelial markers with roles in angiogenesis, endothelial cell cycle regulation, and tissue remodeling, and uncovered depot-specific regulation of Cdkn1c and Efnb1 under cold exposure. These findings underscore the high degree of endothelial specialization in adipose tissue and reveal that BAT-ECs, while quiescent *in vivo*, undergo dedifferentiation and adopt a WAT-like transcriptome when forced into proliferation in culture. Together, this study highlights both the potential and the limitations of cultured adipose-derived ECs, offering a valuable platform to dissect depot-specific endothelial signaling while emphasizing the need to carefully interpret results in the context of the *in vivo* microenvironment and passage number.

## CRediT authorship contribution statement

**Tabea Elschner**: Conceptualization, Data curation, Formal analysis, Investigation, Methodology, Validation, Visualization, Writing – original draft. **Staffan Hildebrand:** Data curation, Formal analysis, Writing – review and editing. **Lara Heubach:** Data curation, Formal analysis, Methodology. **Jana Sander:** Data curation, Formal analysis, Methodology, Writing – original draft. **Stephan Grein:** Data curation, Formal analysis, Resources, Software, Writing – review and editing. **Maria Mircea:** Data curation, Formal analysis, Resources, Software. **Elba Raimundez**: Data curation, Formal analysis, Resources, Software, Writing – review and editing. **Nina Pannwitz:** Data curation, Formal analysis, Methodology. **Anastasia Georgiadi:** Resources, Formal analysis, Writing – review and editing. **Joerg Heeren:** Resources, Writing – review and editing. **Jan Hasenauer:** Supervision, Funding acquisition, Writing – review and editing. **Alexander Pfeifer:** Conceptualization, Funding acquisition, Supervision, Writing – review and editing. **Kerstin Wilhelm-Jüngling:** Conceptualization, Data curation, Formal analysis, Funding acquisition, Investigation, Methodology, Project administration, Resources, Software, Supervision, Validation, Visualization, Writing – original draft, Writing – review and editing.

## Funding

Work here was funded by the Deutsche Forschungsgemeinschaft (DFG, German Research Foundation) – TRR333/1 – 450149205 to T.E., S.H., L.H., S.G., E.R., N.P., J.H., A.P., K.W.J.

## Declaration of competing interest

The authors declare that they have no known competing financial interests or personal relationships that could have appeared to influence the work reported in this paper.

## Acknowledgements

We would like to thank the Core Facility of the Medical Faculty at the University of Bonn for providing support and instrumentation funded by the Deutsche Forschungsgemeinschaft (DFG, German Research Foundation) – Projektnummer 388159768.

## Appendix - Supplementary data

Supplementary Figure 1

Supplementary Figure 2

Supplementary Figure 3

## Data availability

Data will be made available on request.

## Declaration of generative AI and AI-assisted technologies in the manuscript preparation process

During the preparation of this work the author(s) used ChatGPT (*OpenAI*) in order to improve proofreading, readability, and language. After using this tool, the author(s) reviewed and edited the content as needed and take(s) full responsibility for the content of the published article.

**Supplementary Figure 1**

**A** Brightfield image of a pure BAT-EC culture representing a single-layer of cobblestone-shaped endothelial cells.

**B** Representative brightfield image of a fibroblast contaminated BAT-EC culture. Arrowheads depict multi-layered cells with a spiky cell shape, representing fibroblasts. **C** Relative transcript levels of *Pecam1*, *Cdh5*, *Erg* and *Vwf* at four passages of neonatal BAT-ECs (left panel) and WAT-ECs (right panel) and in muMECs, displaying that BAT-ECs and WAT-ECs do not lose endothelial marker expression during the four passages and endothelial transcript levels are higher than in muMECs.

**D** Uniform Manifold Approximation and Projection (UMAP) displaying the different cell types in BAT. All further violine blots display expression levels of the chosen gene within single cells in the depicted cell types:

**E** *Gpd1*

**F** *Pth1r*

**G** *Rgcc*

**H** *Thrsp*

**I** *Cdkn1c*

**J** *Fasn*

**K** *Myo10*

**L** *Chchd10*

**M** *Ccdc85a*

**N** *Efnb1*

**O** *Tbx5*

**P** *Ucp1*

**Q** *Clic5*

**R** *Cdh23*

**S** *Tcf15*

**Supplementary Figure 2**

All UMAP blots display transcript levels measured for the respective genes in single cells isolated from mice house at room temperature (RT; left panel) and house at 5°C for 7 days (cold7; right panel):

**A** *Rgcc*

**B** *Cdkn1c*

**C** *Ccdc85a*

**D** *Tcf15*

**E** Table summarizing log2 fold-change and adjusted p-value of respective genes within the endothelial cell cluster comparing ECs isolated from mice housed at RT versus housed at cold7.

**Supplementary Figure 3**

**A** UMAP displaying the different cell types in BAT. All further violine blots display expression levels of the chosen gene within single cells in the depicted cell types:

**B** *Cfh*

**C** *Col23a1*

**D** *Scn5a*

**E** *Prelp*

**F** *Sned1*

**G** *Smoc1*

**H** *Lrrn4cl*

**I** *Prox1*

**J** *Pcsk6*

**K** *Tll1*

**L** *Loxl2*

**M** *Pard6g*

## Notes

### Competing Interest Statement

The authors have declared no competing interest.

