## Supplementary Figure 1-3 for "From Brown to White: Brown Adipose Tissue Endothelial Cells whiten in Culture Conditions"

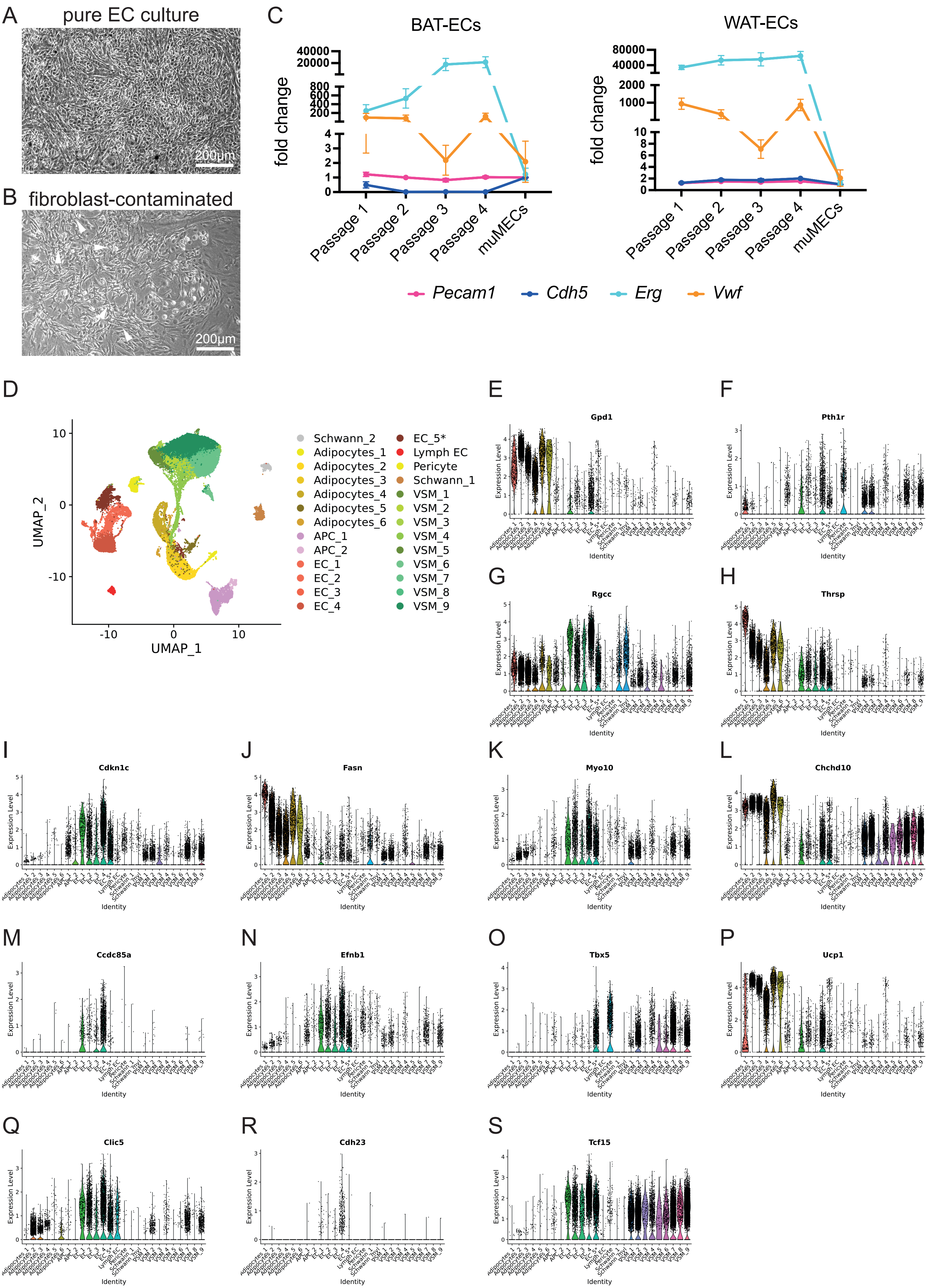

Supplementary Figure 2

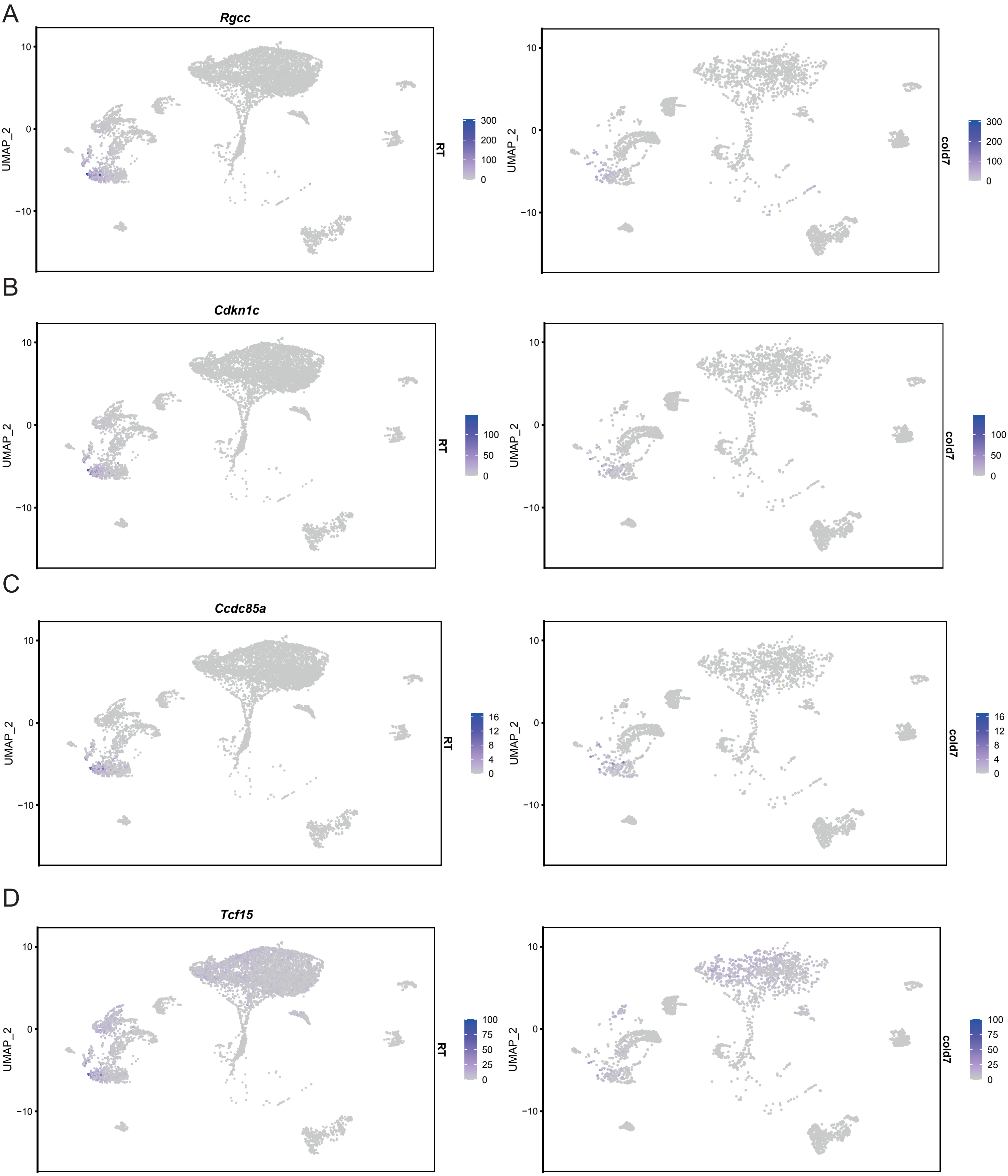


E

| Gene | average log2FC | p-value adj. | Condition |
|---|---|---|---|
| *Rgcc* | 1,9824 | 3,9357E-53 | RT vs cold7 |
| *Cdkn1c* | 1,6348 | 6,0333E-29 | RT vs cold7 |
| *Ccdc85a* | 0,5357 | 1,1301E-13 | RT vs cold7 |
| *Tcf15* | 1,2957 | 2,9378E-31 | RT vs cold7 |
| *Meox2* | 0,4602 | 2,3310E-07 | RT vs cold7 |

### Supplementary Figure 3

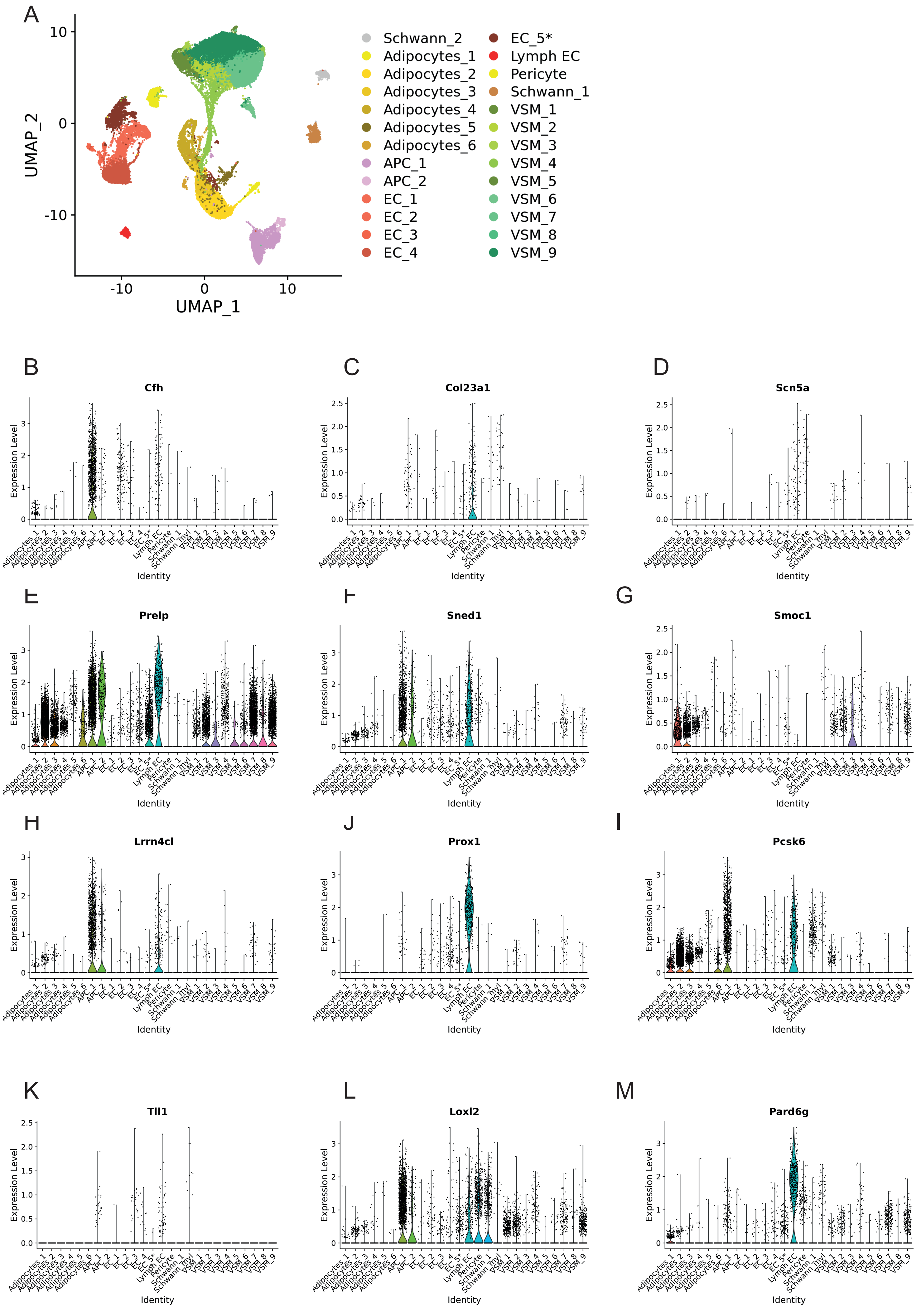
